# Label-free Imaging of the Reversible Rhodopsin Dynamics in a Living Eye

**DOI:** 10.1101/2025.04.02.646910

**Authors:** Yueming Zhuo, Huakun Li, Mohajeet Bhuckory, Davis Pham-Howard, Daniel Palanker

**Author notes:** These authors contributed equally to this work (Yueming Zhuo, Huakun Li).

## Abstract

Vision begins with photoisomerization of a retinal, triggering further conformational changes, followed by a phototransduction cascade in photoreceptors. Microelectrode recordings revealed an early receptor potential (ERP) signal accompanying these conformational transitions. Such techniques are invasive, obscured by multiple electrical processes in the retina, and in rods, they are limited to the contribution of a small fraction of rhodopsin embedded in the nascent disc membrane and plasma membrane. Recent advances in phase-sensitive OCT (Optoretinography, ORG) enable detection of nanoscale deformations of retinal cells associated with physiological processes. However, in previous ORG studies, focused primarily on cones, cell deformation related to ERP was largely obscured by osmotic swelling and long stimuli did not resolve the isomerization dynamics of photopigments.

Here, we demonstrate a very robust electromechanical signature of photoisomerization in rods at microsecond-scale temporal resolution. Green flash induces sub-millisecond-fast contraction of the outer segments by hundreds of nanometers, while subsequent UV flash reverts the activated molecules, producing an opposite response of a similar magnitude. This approach surpasses the sensitivity of electrical methods by integrating the response across all the discs in rod outer segments. Noninvasive and label-free imaging of the rhodopsin dynamics in a living eye opens the door to fundamental studies of visual transduction and to differential diagnosis of the photoreceptors’ dysfunction.

## I. Introduction

Vision begins with the absorption of photons in photoreceptors, triggering photoisomerization of photopigments and subsequent phototransduction, ultimately hyperpolarizing the cells and modulating their neurotransmitter release [1]. After photon capture, rhodopsin undergoes a series of isomerization intermediates, including lumirhodopsin and metarhodopsin I (Meta I), before reaching the biochemically active metarhodopsin II (Meta II) state [2, 3], a process involving charge transfer across the membrane, primarily in the outer segment discs [4–6]. Patch-clamp electrophysiological recordings in photoreceptors, a very invasive technique applicable only ex vivo, revealed a rapid electrical signal, called the early receptor potential (ERP) [6–8]: a brief depolarizing R1 phase corresponding to the formation of Meta I, and a dominant hyperpolarizing R2 phase reflecting the conversion from Meta I to Meta II [8]. A UV flash (400 nm) delivered to photoreceptors fully bleached by green light can elicit a rapid electrical signal of opposite polarity, indicating that UV photons can revert the activated rhodopsin into another isoform [9, 10], subsequently shown to be Metarhodopsin III (Meta III), which can again absorb green photons [11].

In vivo observation of these processes is critical for understanding the biophysics of vision in real physiological conditions and for differential diagnosis of multiple forms of photoreceptors’ disfunction. Indeed, reduction in ERP amplitude has been noted in patients with diabetic retinopathy and retinitis pigmentosa [12]. The electrical signal accompanying photoisomerization originates in charge transfer across the cell membrane [5, 6]. Even though rods represent a vast majority of photoreceptors in most mammalian retinas [13], such recordings from rods are more challenging than from cones since most of rhodopsin is embedded in the sealed disc membranes, which are isolated and electrically decoupled from the plasma membrane of the outer segment (Fig. 4A) [3, 14], unlike cones, where all the discs are folded into a common plasma membrane itself [15]. As a result, rod signals are overshadowed by cone responses in primate and human retinas [16, 17].

Phase-sensitive optical coherence tomography (OCT) offers a non-invasive approach to detecting cellular processes by measuring nanometer-scale deformations in photoreceptors associated with metabolic events, including photoisomerization and phototransduction [18–24]. This technique, termed Optoretinography (ORG), enabled observation of a rapid (few milliseconds) contraction and slow (hundreds of milliseconds) expansion of the cone outer segments (COS) immediately following a light stimulus [18–20, 25]. Contractile deformations were attributed to electromechanical coupling [26] - changes in the surface tension induced by charge transfer across the membrane, while the slower expansion of COS - to osmotic influx of water during phototransduction, as well as the swelling of the cone opsin and disc membrane [25, 27].

Typically, the swelling in cones obscures the rapid contractile signal related to ERP, and relatively long light stimuli used in earlier ORG studies [18–20] could not resolve the sub-ms dynamics of the rhodopsin isomerization [2, 3, 10].

By employing ultrafast (10-kHz B-scan rate) OCT recordings and microsecond-scale light stimuli, we observed a sub-ms fast contraction of the rod outer segments (ROS) by hundreds of nanometers that persisted for hundreds of milliseconds. About a thousand ROS disc membranes collectively contributed to this robust electromechanical effect, surpassing the sensitivity of electrical measurements. Furthermore, a UV (385 nm) flash delivered after extensive bleaching by green (520 nm) light reversed the contraction and enabled further absorption of green photons, indicating a transition from Meta II to Meta III. This imaging method with microsecond-scale temporal resolution opens the door to fundamental studies of visual transduction in vivo and differential diagnosis of the photoreceptors’ dysfunction, with a possibility of single-cell spatial resolution.

## II. Results

### A. Dynamics of the ROS contraction

To assess the dynamics of the ROS contraction, we applied green (520 nm) flashes to retinas of wild-type rats while acquiring repeated B-scans at a 10 kHz frame rate using a custom-built ultrafast line-scan OCT system. To investigate the impact of pulse duration on the waveform of the rapid contractile response (Fig. 1), we employed pulse durations of 50, 100, 200, 500 µs and 1 ms (see Fig. S4 for temporal profiles) and adjusted the power to maintain a consistent pulse energy - 50.6 *±* 0.2 µJ, measured in front of the cornea. We also explored the stimulus strength-dependence of the rapid contraction by fixing the pulse duration at 100 µs and adjusting the flash energy in approximately two-fold increments (Fig. 2).

**Fig. 1.**
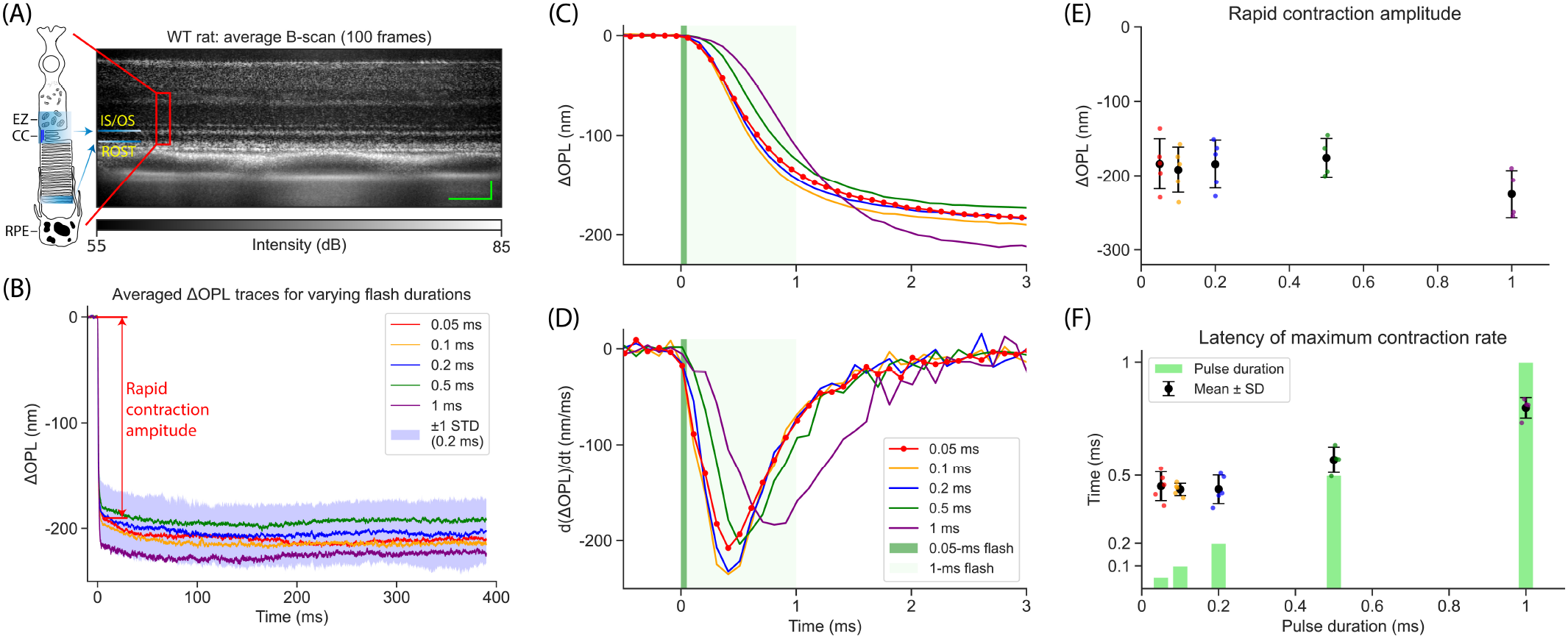
Dynamics of the ROS contraction evoked by flashes with varying pulse durations. (A) Representative structural image of a wild-type (WT) rat retina, with an inset illustrating two retinal bands – IS/OS and ROST – utilized for calculating the optical path length (OPL) along the rod outer segment (ROS). Scale bars: 50 µm. IS/OS: inner segment/outer segment junction; ROST: rod outer segment tips; EZ: ellipsoid zone; CC: connecting cilium; RPE: retinal pigment epithelium. (B) ΔOPL traces extracted from ROS and averaged across 4-5 animals for each flash duration. The purple band represents the standard deviation range for 0.2 ms pulse duration. The rapid contraction amplitude is extracted by averaging ΔOPL within the 3-4 ms time window. (C) Enlarged view of ΔOPL traces within the first 3 ms. (D) Time derivatives of ΔOPL traces. (E) Amplitude of the ROS rapid contraction. Scatter plot of all individual data points, with error bars indicating the standard deviation range for each pulse duration. (F) Latency of the maximum contraction rate. Statistical distribution of the time points corresponding to the maximum contraction rate in various animals for each pulse duration, with error bars representing the standard deviation range. Green bars represent varying pulse durations for comparison.

**Fig. 2.**
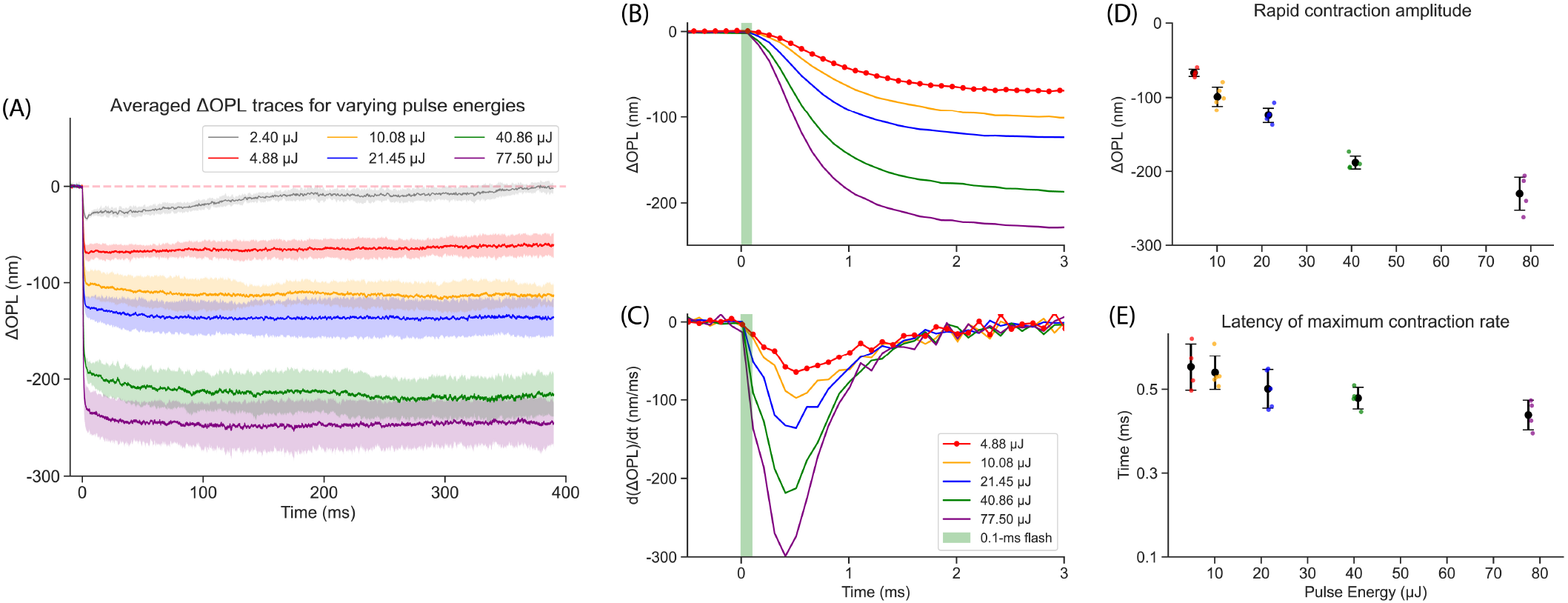
ROS contraction with varying pulse energies. (A) ΔOPL traces in response to 100-µs pulses with varying energies. Semitransparent band represents the standard deviation range. (B) ΔOPL traces within the first 3 ms after the flash. (C) Contraction rate – the time derivative of ΔOPL traces. (E) Amplitude of the rapid contraction: scatter plot of all individual data points, with error bars indicating the standard deviation range for each pulse energy. (F) Latency of the maximum contraction rate: data points of individual animals, with error bars representing the standard deviation range.

Changes in the phase difference between two retinal bands reflect variations in the optical path length (ΔOPL) between them. As illustrated in Fig. 1A, to assess the light-evoked response of the ROS, we computed the phase difference between the inner segment/outer segment (IS/OS) junction and the rod outer segment tips (ROST, the top of the thick hyperreflective band consisting of ROST and the retinal pigment epithelium). The ΔOPL traces presented in Fig. 1B for each pulse duration were extracted from ROS and averaged across 4-5 rats. Since similar photon densities were delivered to the retina, the resulting rapid contraction amplitudes of the outer segments were comparable, although response to the 1-ms flash was noticeably slower and its contraction amplitude was slightly larger than those with shorter pulses. This is likely due to a more pronounced photo-reversal effect of shorter pulses, resulting in fewer activated rhodopsins (see section Modeling the ROS contraction). Amplitudes of the rapid contraction were extracted by averaging ΔOPL within the first 3-4 ms temporal window and their statistical distributions are shown in Fig. 1E.

In stark contrast to the fast recovery (*<*20 ms) from the rapid contraction observed in COS responses [20], the ROS remained in a contracted state throughout the 400-ms acquisition window. As can be seen in Fig. 1C, the contraction process begins almost immediately after the flash onset, but continues for about 3 ms, even with the 50-µs flash. The contraction rate, shown in Fig. 1D, was calculated by taking a time derivative of each trace. Latency of the maximum contraction rate was extracted and compared to pulse durations in Fig. 1F. Interestingly, for flashes shorter than 0.5 ms, latency of the maximum contraction rate remains almost the same - about 0.45 ms, and it starts increasing for longer pulses.

While a short, intense flash revealed the impulse response of the rapid contraction process, its strength dependence provides additional insights into the underlying mechanisms. Fig. 2A displays the ΔOPL traces in response to 100-µs flashes with varying energy levels. As expected, amplitude of the rapid contraction increases with incident photon density. With the lowest pulse energy, OPL significantly recovered during the 400 ms after the flash, while at higher energy levels, contraction remained steady and even slowly increased over time. Dynamics of the rapid contraction within the first 3 ms is shown in Fig. 2B. The contraction amplitudes, determined by the same procedure as in Fig. 1, are shown in Fig. 2D, revealing a saturating trend. Latency of the maximum contraction rate, shown in Figs. 2C and 2E, decreases with increasing pulse energy from about 0.55 ms and saturates at around 0.42 ms.

### B. Modeling the ROS contraction

#### Contraction amplitude as a function of bleach level

Early photoproducts, up to Meta I, can absorb a second photon and restore either rhodopsin or isorhodopsin, a process called photo-reversal [28–30]. Therefore, if the flash duration is shorter than the thermal decay time constant from Meta I to Meta II (a few hundred microseconds in rodents [5, 10]), photoreversal of rhodopsin bleaching cannot be neglected. To determine the bleach levels induced by microsecond-scale flashes in our experiments, we considered all critical light-driven and thermal processes involved in flash photolysis (Fig. 3A and Supplementary Information: Bleach level calculation). The calculations showed that rapid contraction amplitude ΔOPL increased logarithmically with the estimated bleach levels (Fig. 3B).

**Fig. 3.**
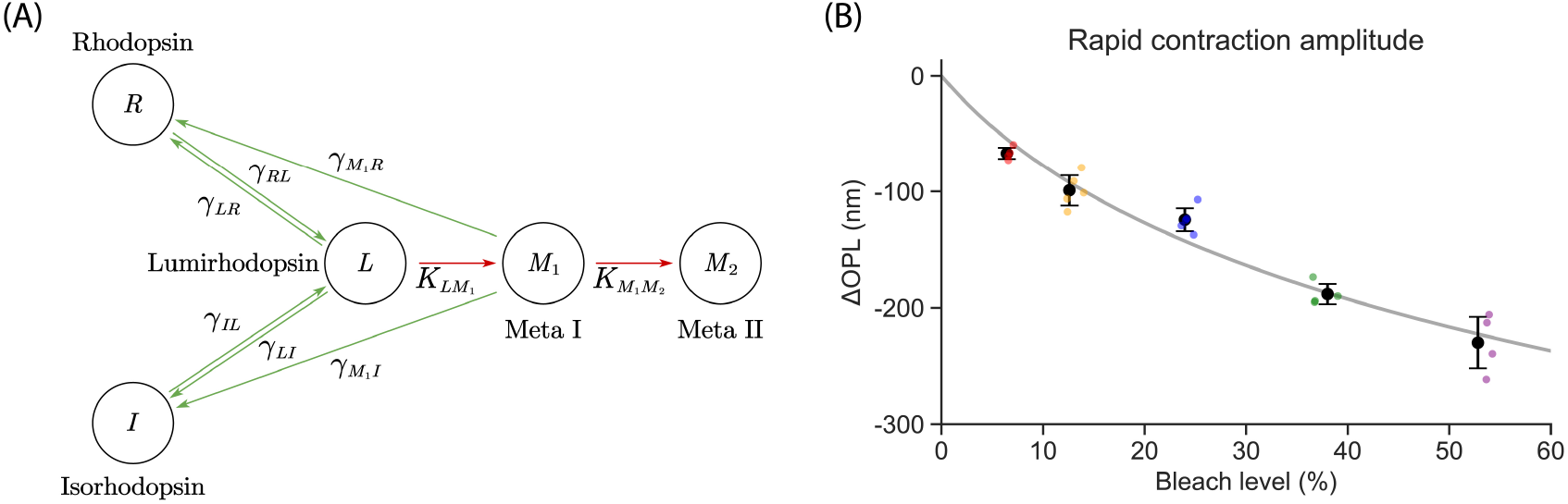
Rhodopsin isomerization pathways in flash photolysis and the dependence of rapid contraction amplitude on bleach levels. (A) Rhodopsin isomerization pathways include photon-driven transitions (green arrows) and thermal decay processes (red arrows). *γ*_*XY*_ : quantum efficiency of the photoisomerization from X to Y. *K*_*XY*_ : reaction rate of the thermal decay from X to Y. (B) Rapid contraction amplitude, measured in Fig. 2D, increased logarithmically with the estimated bleach level. Colored dots represent individual measurements, and the gray line denotes the logarithmic fitting.

#### Modeling the mechanical deformation of OS

The rapid contraction measured in COS was previously explained using the voltage-dependent membrane tension model [26]. In brief, the charge shift during the R2 phase of the ERP leads to hyperpolarization of disc membranes. Increase of charge density in the Debye layer increases the repulsion of ions [31– 33], resulting in lateral stretching of the disc membrane. Due to conservation of the discs’ volume during millisecond-scale dynamics, this lateral expansion leads to the axial contraction of the discs. As detailed in Supplementary Information, these considerations yield a good match between the modeled and measured contraction amplitude across varying bleach levels (see Fig. S1). Furthermore, the values of the fitted parameters are comparable to typical values in the literature, suggesting that the observed contraction aligns with the voltage-dependent membrane tension model.

However, this quasistatic model failed to capture the latency of the contraction (see blue curves in Figs. 4B-C), indicating a potential involvement of viscous effects. One factor that could dampen the response, as proposed in our previous study [26], is the viscous force exerted by surrounding fluid when individual discs expand laterally. Another mechanism, proposed in this study, arises from the friction when discs move axially in the viscous cytoplasm. Axial shrinkage of each disc results in stretching of the spacers connecting adjacent discs [27, 34, 35], which then drive axially the discs and ROST. The viscous force induced by the surrounding cytoplasm during this axial movement dampens the contraction process.

**Fig. 4.**
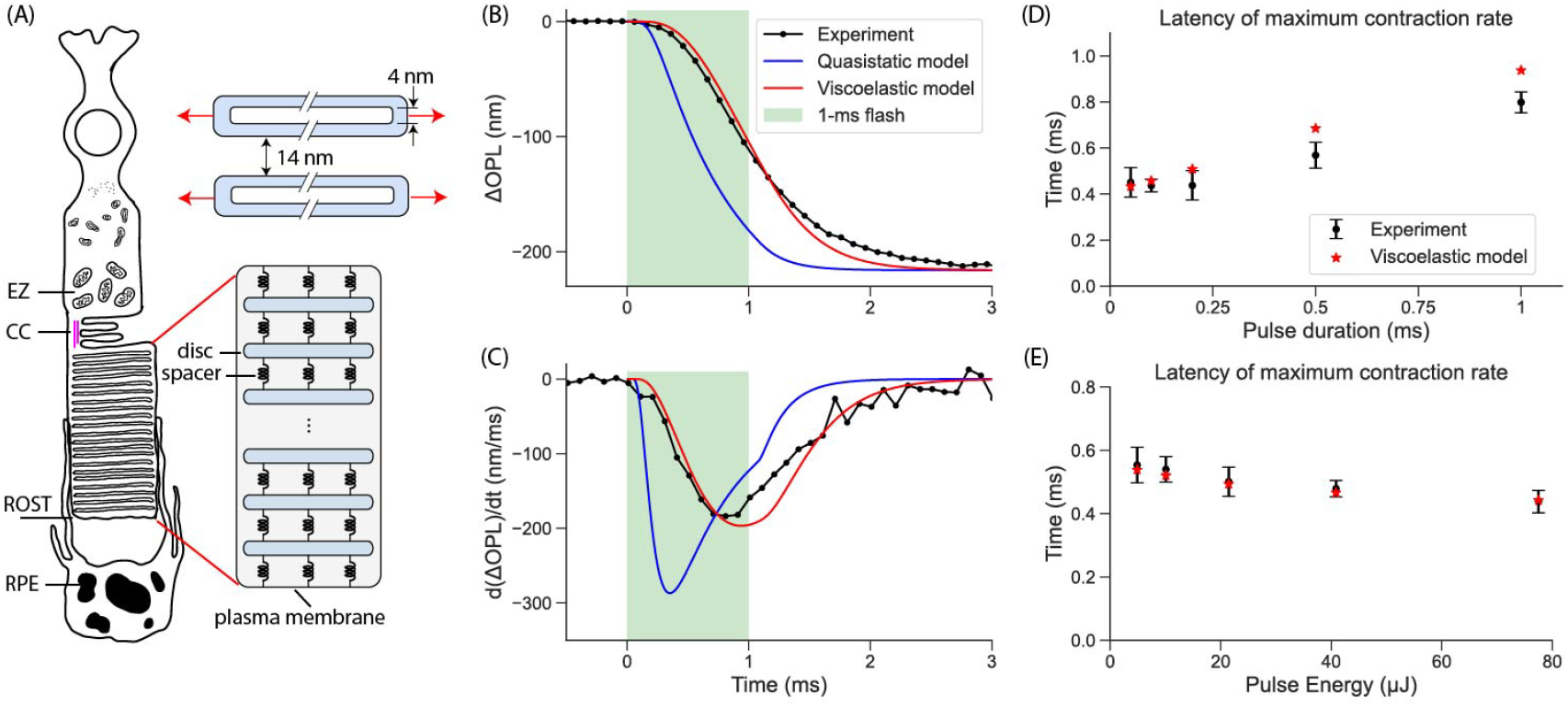
Viscoelastic modeling of the rapid ROS contraction. (A) Diagram of a rod photoreceptor. The top right inset illustrates lateral disc expansion upon light absorption. The bottom inset depicts the inter-disc spacers connecting the discs. (B) Experimentally measured ΔOPL response to a 1-ms flash and predictions of the quasistatic model and the planar & axial viscoelastic model. (C) Contraction rate obtained by time derivative of the traces in (B). (D) The best-fit model and experimental data for latency of the maximum contraction rate with varying pulse durations. (E) Similar comparison as in (D) for varying pulse energies.

A combined model considering both, the lateral and axial viscous and elastic forces applied to each disc, enabled a better fit to experimental results (Fig. 4 and Supplementary Information: Computational Model for ROS Contraction). As shown in Figs. 4B and C, using the 1-ms pulse measurement as an example, the model provides a better fit for the temporal evolution of ΔOPL than its quasistatic approximation. The best fit yielded damping coefficients of 4.0 *×* 10^5^ N · s · m^−3^ for the lateral expansion of disc membranes and 4.4 *×* 10^4^ N · s · m^−3^ for the axial movement of the discs. These damping coefficients correspond to the linear damping behavior caused by cytoplasmic sheets with thickness of approximately 10 nm and 100 nm, respectively [36], which is reasonable compared to the membranous structure of ROS [34, 35]. As shown in Fig. 4D and 4E, the model-derived latency of the maximum contraction for varying pulse durations and energy levels matches the experimental data much better than the quasistatic model.

### C. Reversed isomerization with UV flash

A UV flash delivered to Meta II can elicit an electrical signal of opposite polarity compared to ERP [9, 10]. To assess the ORG response related to this process, a 1-ms flash at 385 nm was introduced after the rhodopsin bleaching by a 1-ms green flash. Pulse energy of the green flash was sufficiently high (340 µJ) to bleach nearly all rhodopsin, as validated by the very small residual response to the second green flash 200 ms later (Fig. 5A). At this pulse energy, thermal effects were still negligible compared to ORG signals (see Discussion).

**Fig. 5.**
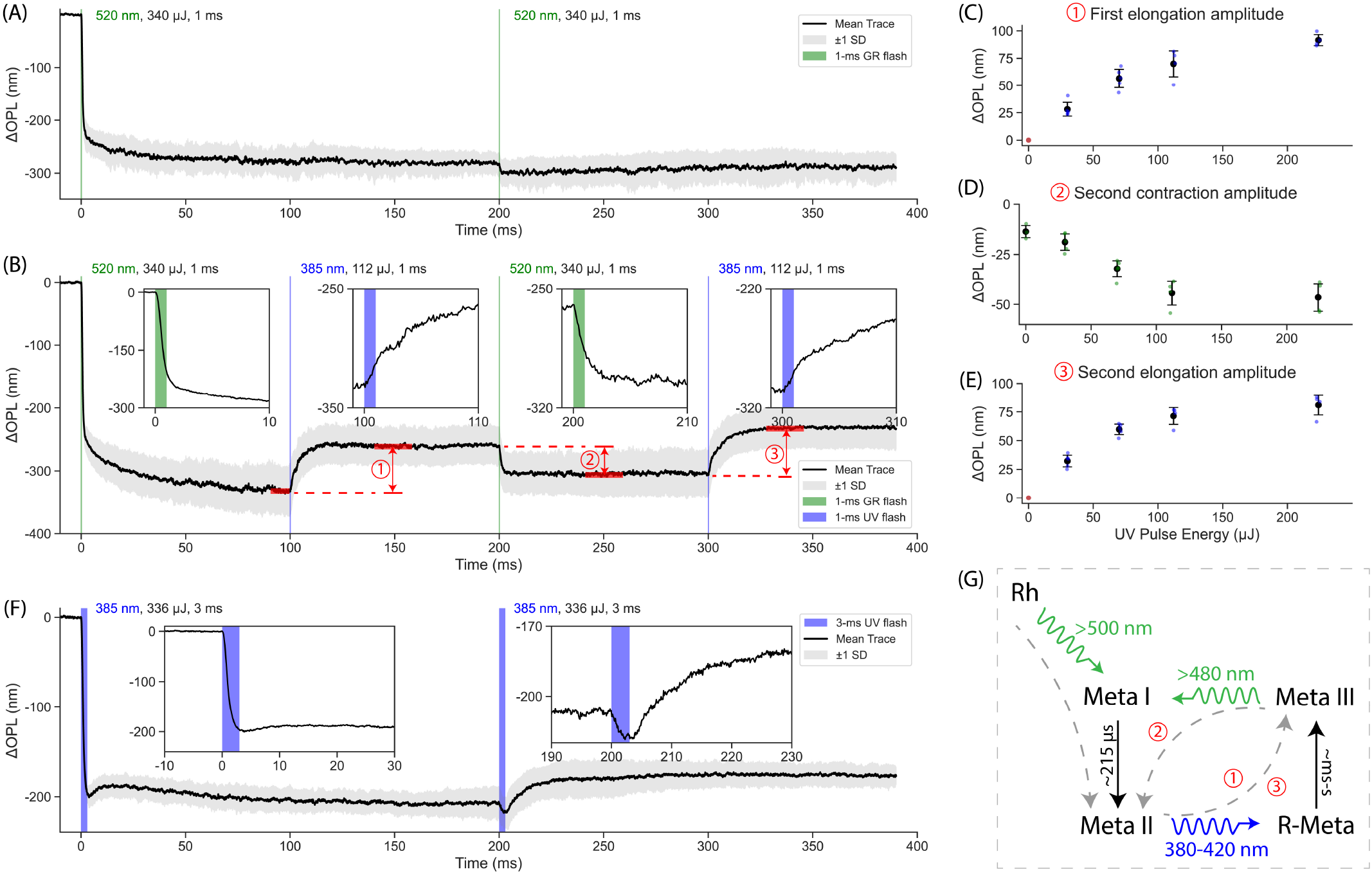
Reversible isomerization with green and UV stimulation. (A) Control experiment: ORG response elicited by two green flashes (1 ms, 336 µJ, 200-ms apart). (B) Response to a flash sequence of green-UV-green-UV, with a UV flash of 1 ms, 112 µJ. Insets show the first 10-ms time window. Red arrows indicate the amplitudes plotted in panels C-E. (C) The first elongation amplitude as a function of UV pulse energy. Dots show measurements in individual rats, error bars – one standard deviation around the average. (D) The second contraction amplitude as a function of UV pulse energy. (E) The second elongation amplitude as a function of UV pulse energy. (F) ORG responses to two UV pulses (3 ms, 336 µJ, 200-ms apart). Insets show the first 30-ms time window. (G) Rhodopsin isomerization pathways related to this study (more detailed diagram in Supplementary Information, Figs. S8 and S9).

UV flashes were delivered 100 ms after each green pulse at UV energy levels of 30, 70, 112 and 224 µJ. As shown in Fig. 5B, the first UV flash (1 ms, 112 µJ), applied to an extensively bleached retina, elicited a pronounced elongation of the ROS (with ∼10 ms time constant), suggesting a reversed response relative to the contraction induced by the first green flash. The second green flash, applied 100 ms after the UV pulse, induced stronger contraction compared to the control (Fig. 5A), implying partial recovery of photopigment, restoring its ability to respond to green photons. The final UV flash elicited another elongation, similar in amplitude to the first UV pulse.

Amplitude of the elongation produced by the first UV flash exhibited a saturating trend with pulse energy, as shown in Fig. 5C. As anticipated, higher energies of the first UV flash increased the contraction amplitude for the second green flash (Fig. 5D) since more photopigments sensitive to green photons were regenerated by more intense UV light, although this effect saturated beyond 112 µJ. Amplitude of the elongation induced by the final UV flash followed a pattern similar to that of the first one (Fig. 5E).

Remarkably, when UV flash was applied to a dark-adapted retina alone, without a preceding green pulse, it elicited a completely different ORG response (Fig. 5F) - rapid contraction by 200 nm, similar to the response observed with a green flash. After the rapid contraction, however, the ROS briefly expanded by a few tens of nm, before returning to its contracted state. The second UV flash 200 ms later produced a small additional contraction (like the second green flash in Fig. 5A) but followed by a more pronounced elongation (∼20 ms time constant), lasting for hundreds of ms.

## III. Discussion

### A. ΔOPL due to refractive index variations and thermal effects

The measured ΔOPL in ROS can stem from a variation in the refractive index, a mechanical deformation, or a combination of both. To quantify the refractive index change due to isomerization of photopigments, the Kramers–Kronig transform was applied to their absorption spectra. Following absorption of a green photon, rhodopsin is converted to Meta II, leading to a pronounced shift of its absorption peak from 500 nm to 380 nm [3, 37, 38], as shown in Fig. S5A. Assuming the other peaks in UV range do not change, the resulting refractive index change at the OCT center wavelength (840 nm) is on the order of 10^−5^, corresponding to ΔOPL *<* 1 nm along the ROS (Fig. S5B).

Light absorption in pigmented retinal layers, such as photoreceptor OS, retinal pigment epithelium and pigmented choroid, also results in heating, which can change refractive index and induce thermal deformations [39]. Our thermomechanical model [40] shows that the dominant effect is thermal expansion of the retinal layers. At the highest bleach level in our experiments, temperature rise is *<* 0.5 ^°^C and the associated thermal expansion of the outer segments is *<* 10 nm (Fig. S6). This thermal effect is of the opposite sign and of a negligible amplitude compared to hundreds of nanometers of the ROS contraction observed in our experiments.

### B. ORG in cones and rods

In COS, a rapid contraction (*<*20 ms) is followed by hundreds of ms-long expansion [20], which has been attributed to osmolarity-driven water influx, as well as the swelling of cone opsins and disc membranes during phototransduction [25, 27]. Amplitude of both the rapid contraction and slow expansion increase with light intensity [20]. Like the temporal overlap between ERP and late receptor potential (LRP) in electrical recordings [7, 8], the expansion phase in cone ORG obscures part of the contraction.

In rod ORG, expansion is much weaker and slower, and hence it does not obscure the contraction. This could contribute to the significantly larger contraction amplitude observed in rat ROS (*>* 200 nm, Fig. 2D) compared to that in human COS (∼50 nm [20]). In fact, slow expansion was observed only at very low bleach levels (*<* 3%), as can be seen in Fig. 2A and Fig. S7. At higher bleach levels, ROS remains contracted for at least 400 ms. One reason for small expansion could be structural: except for a few nascent disc membranes, ROS is primarily composed of densely packed, sealed discs enclosed within a tight plasma membrane [3, 14], which has very limited room for expansion, unlike COS, where the discs are infoldings of the common plasma membrane, providing plenty of room for expansion [15]. Lack of expansion and even an additional contraction during the first 100 ms at higher bleach levels in ROS (Fig. 1B and 2A) indicate another potential contraction mechanism, which counterbalances the osmotic swelling at higher bleach levels.

The remarkable difference in membrane time constants between rod and cone disc membranes may also contribute to the much slower recovery in ROS. Patch clamp-recorded ERP recovers within a few ms due to the passive discharge through the plasma membrane, governed by its resistance and capacitance [7, 8]. The time constant of Meta II formation in human M-cone visual pigment, which induces the hyperpolarizing R2 phase of the ERP, is approximately 6 ms [41]. This value is on the same scale as the typical membrane time constant [8], indicating that a passive discharge of the disc membrane in COS reduces the transmembrane potential change and the associated contraction amplitude. However, unlike plasma membrane, rod disc membrane has exceptionally high specific membrane resistance (on the order of MΩ· cm^2^) due to the absence of ion channels [42, 43], yielding a much larger membrane time constant - on the order of seconds (with a typical specific membrane capacitance of 1 µF ·cm^−2^) [6]. Consequently, the very slow passive discharge of the disc membrane in rods maintains the hyperpolarization induced by the initial photoisomerization, as well as the associated mechanical contraction, for a long time.

### C. Rod ORG responses to UV

Ex vivo electrical recordings demonstrated that an intense UV (400 nm) stimulus applied to a fully bleached retina induces electrical response opposite in polarity to the ERP [9, 10]. Moreover, a second green flash following the UV pulse elicits an ERP response with the same shape as that of the pre-bleached retina (albeit with a lower amplitude), suggesting that photopigment was regenerated by the UV flash. Diagram of the photoisomerization and thermal relaxation pathways in rhodopsin shown in Fig. 5G and Fig. S8 suggests that UV light (380-420 nm) can trigger 15-anti/15-syn isomerization in Meta II, leading to formation of reverted-Meta (R-Meta), which is followed by Schiff base protonation, resulting in formation of Meta III [3, 37]. When illuminated by green light, Meta III undergoes a 15-syn/15-anti isomerization around the Schiff base C=N bond, reverting to Meta I, which is then followed by Schiff base deprotonation, forming Meta II again. Hence, the Meta III photointermediate, though structurally different from the ground state rhodopsin, can absorb green light and induce similar state transitions (Fig. 5G and Fig. S8). We conjecture that the rapid (sub-ms) contraction of rod discs arises from the transition of Meta I to Meta II via Schiff base protonation (in hundreds of microseconds in rodents) [5, 10], while the reverse process - transition from R-Meta to Meta III via Schiff base re-protonation (∼ms to s in various in vitro preparations) [38] - induces slower (∼10 ms) ROS elongation.

As shown in Fig. 5F, when a UV flash is applied to a dark-adapted retina, the ROS undergoes a rapid contraction, similar to that observed with a green stimulus (Fig. 5A). This finding indicates that UV photons can also trigger the transition from rhodopsin to Meta II. As a result, prior to the second UV flash, the ROS harbors a mixture of rhodopsin, Meta I, Meta II and a small fraction of Meta III. Upon the second UV flash, three concurrent reaction pathways can be activated: (i) rhodopsin transitions through Meta I to Meta II, (ii) Meta II transitions to Meta III, and (iii) Meta III transitions back to Meta II through Meta I (see Fig. S9). The first and third pathways induce the ROS contraction, while the second pathway elicits elongation. The dynamic interplay of these processes governs the morphological response of the ROS in Fig. 5F.

### D. ORG vs. ERG: translation to clinic

ERG is widely used in basic studies of retinal physiology and for ophthalmic diagnostics [44]. ERP, which reflects the pigment isomerization processes, can provide information about the visual pigment mutations and visual retinoid cycle. The following slower component of the LRP or a-wave helps diagnose the issues associated with phototransduction. ERP amplitude in rods is much smaller than in cones because this signal reflects the charge shift across the plasma membrane, where ROS contains significantly fewer visual pigments than in the isolated disc membranes enclosed within the cell membrane. In contrast, optical signature of photoisomerization in rods is much stronger than in cones and provides a very convenient label-free non-invasive alternative to ERG for retinal diagnostics. Due to cumulative motion of the isolated discs in rods, optical approach is far more sensitive than electrical recordings, reducing the need for multiple intense stimuli, and providing functional information, co-registered with structural OCT imaging, potentially down to single cell resolution when combined with adaptive optics [19, 21, 22].

## IV. Materials and Methods

### A. Phase-sensitive OCT imaging

The high-speed line-scan spectral-domain OCT was assembled according to the optical layout shown in Fig. 6. The OCT beam from the supercontinuum laser (FIR-9, NKT Photonics, Denmark) was collimated using lens CL1 (AC254-060-B, Thorlabs) and filtered through two spectral filters (Semrock: FF01-776/LP-25, Thorlabs: FESH0900), producing a bandwidth with a full width at half maximum (FWHM) of about 120 nm centered at 840 nm.

**Fig. 6.**
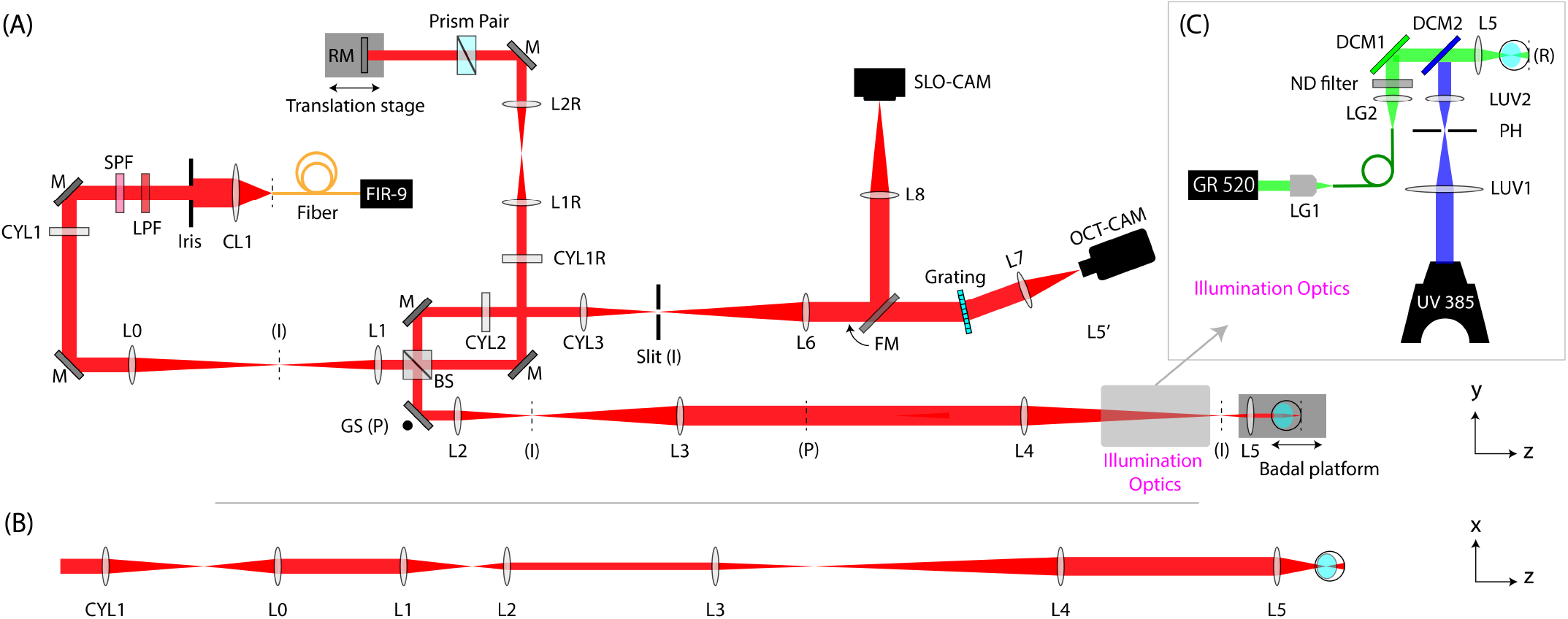
The optical setup. (A) Top view (y-z plane) of the setup. CL: collimating lens. SPF: short pass filter. LPF: long pass filter. CYL: achromatic cylindrical lens doublet. L: achromatic lens doublet. M: mirror. RM: reference mirror. BS: beamsplitter. GS: galvo scanner. DCM: dichroic mirror. FM: flippable mirror. PH: pinhole. DPSS: diode-pumped solid-state laser. FIR-9: NKT FIR-9 OCT laser. The effective focal lengths: CL1=60 mm, CYL1=CYL1R=CYL2=L4=L8=250 mm, L0=150 mm, L1=L1H=L7=100 mm, L2=CYL3=75 mm, L3=L1R=L2R=L2H=L6=200 mm, CL2=L5=30 mm. (I): conjugate image planes. (P): conjugate pupil planes. Not to scale. (B) Unfolded x-z view of the OCT illumination path showing line-field on the retina. (C) Optical paths for green (520 nm) and UV (385 nm) stimulation. DCM1-2: Dichroic mirror 1 and 2. R: retinal plane. The effective foal lengths: LG2=30 mm, LUV1=100 mm, LUV2=35 mm.

The beam then passes through a cylindrical lens (CYL1) which forms a line image along the y-axis at the focal plane. Following optical conjugation through the afocal telescope L0-L1, the beam was divided into reference and sample arms using a non-polarizing beamsplitter (BSS11, Thorlabs) with a 30:70 (R:T) ratio. Two more afocal telescopes in the sample arm, L2-L3 and L4-L5, conjugates the one-dimensional galvo scanning (GS) mirror (8310K Series Galvanometer, Cambridge Technology) to the system pupil plane. The lens L5 and the animal fixation stage were both placed on the Badal platform to adjust beam focus on the retina. The elliptical reference beam after the beamsplitter was re-collimated by a cylindrical lens (CYL1R), followed by an afocal telescope (L1R-L2R). A prism pair (#43-649, Littrow Dispersion Prism, Edmund Optics) was used to balance the dispersion mismatch between the reference and sample arms. In the detection path, an anamorphic telescope (L2-CYL2-CYL3) was used to conjugate image planes for optimizing the spatial and spectral resolution simultaneously. An adjustable slit was placed at the image plane to minimize stray light or back-reflections from the contact lens.

The backscattered sample beam and reflected reference beam were diffracted by a 600 l/mm grating (WP-600/840-35 *×* 45, Wasatch Photonics) and the image was finally conjugated to the detector of the high-speed camera (Phantom v641) via the telescope (L6-L7), yielding a raw image size of 768 *×* 512 pixels (spectral *×* spatial). In this arrangement, each frame of the camera generates a B-scan of the retina. The scanning laser ophthalmoscope (SLO) camera was aligned to be parfocal with the OCT camera. SLO imaging was used for finding a focal plane of the retina, while adjusting the Badal platform, prior to OCT imaging.

### B. Optical paths of retinal stimulation

The green flash path was coupled with the OCT illumination path using a dichroic mirror (NFD01-532-25 *×* 36, Semrock). The output of the green laser (520 nm semiconductor laser, Civillaser) was coupled into an optical fiber (MHP200L02 - Ø200 µm Core, 0.22 NA, Thorlabs) using an objective lens LG1 (Fig. 6C). The output of the fiber was optically conjugated to the system pupil plane via an afocal telescope (LG2-L5). The illuminated area on the retina was estimated to be 1.88 mm^2^, based on a standard rat eye model [45].

For UV stimulus, the collimated output of a high-power LED (385nm, SOLIS-385C, Thorlabs) was focused by LUV1 and illuminated a 2.5-mm pinhole (P2500K, Thorlabs) at an intermediate image plane, which was conjugated to the system pupil plane using an afocal telescope (LUV2-L5). At the pupil plane, the beam size was about 2 mm, ensuring that the beam will not be cropped by the rat pupil on its way to the retina. The illuminated area on the retina was estimated to be 1.56 mm^2^. The UV light was coaxially aligned with the OCT beam using DCM2 (FF495-Di03-25*×*36, Semrock).

### C. Animal preparation

All experimental procedures were approved by the Stanford Administrative Panel on Laboratory Animal Care and conducted in accordance with the institutional guidelines and conformed to the Statement for the Use of Animals in Ophthalmic and Vision research of the Association for Research in Vision and Ophthalmology (ARVO). Long Evans rats (N = 15) were used in this study, and all experiments were performed when they were between P75 and P120 of age. Animal colonies were maintained at the Stanford Animal Facility in 12-hour light/dark cycles with food and water ad libitum.

For all the ORG experiments, animals were dark-adapted more than 12 hours overnight. Animals were anesthetized with a mixture of ketamine (75 mg/kg) and xylazine (5 mg/kg) injected intraperitoneally. The pupils were dilated with a mixture of 2.5% Phenylephrine Hydrochloride and 0.5% Tropicamide (Bausch & Lomb, Rochester, NY) ophthalmic solution and a zero-power contact lens (base curvature 3.00 mm, diameter, 6.00 mm, optical power 0.00 D; Lakewood, CO 80226) was placed on the eye for imaging. To minimize motion artifacts caused by respiration and heartbeat, rat’s head was stabilized using a custom-built fixation stage equipped with a bite bar and ear bars.

### D. Data processing

The complex-valued OCT signals were reconstructed following the conventional processing pipeline, i.e., k-linearization and discrete Fourier transform of the spectral interferogram captured by the camera. Subpixel-level bulk displacements between the first and subsequent B-scans were estimated by locating the peak of upsampled cross-correlation maps [46], then each B-scan was registered to the first B-scan using our custom image registration algorithm [47]. The registered B-scans were flattened along the IS/OS band.

To extract ORG signals, we first calculated the temporal change of phase signals by computing the multiplication of each B-scan with the complex conjugate of the first B-scan,

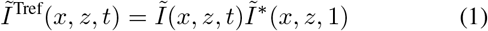

where 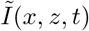 represents the complex-valued OCT images after registration and flattening. *x, z* and *t* denote indices along the lateral, axial and temporal dimensions, respectively. 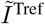 is the time referenced complex-valued OCT signal.

The residual bulk phase errors were cancelled out by calculating phase difference between two retinal layers. For each pixel in the target layer, denoted by (*x*_tar_, *z*_tar_), we selected a reference region from the reference layer - IS/OS in this study. The reference region was centered at *x*_tar_ and spanned across adjacent 11 A-lines. The temporal phase change in the reference region, φ_ref_(*t*), can be calculated by,

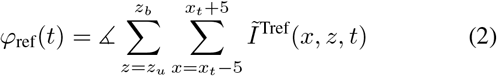

where *z*_*u*_ and *z*_*b*_ denote the upper and bottom boundaries of the reference layer, ∡ denotes the argument operator.

Finally, ORG signals can be obtained by

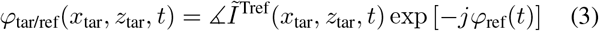

where *j* is the imaginary unit. We then applied the same process to every pixel in the target layer and calculated the average of extracted phase traces across different pixels, yielding φ_tar/ref_(*t*). The phase signal can be converted into changes in OPL by,

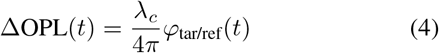

where *λ*_*c*_ is the center wavelength of OCT imaging beam (840 nm).

## Acknowledgements

We thank Dr. David Veysset for fruitful discussions of the ROS modeling. This work was funded by the National Institutes of Health (U01 EY032055), Air Force Office of Scientific Research (FA9550-20-1-0186) and Research to Prevent Blindness.

## Data availability

All data needed to evaluate the conclusions in the paper are present in the paper. Additional data related to this paper may be requested from the authors.

## Author contributions

Y.Z., H.L. and D.P. designed research; Y.Z. and H.L. built the optical setup; Y.Z., M.B. and D.P.H. performed experiments; Y.Z. and H.L. processed the experimental data; H.L. and Y.Z. performed computational modeling; Y.Z., H.L. and D.P. wrote the paper.

