## Supplementary Information for "Label-free Imaging of the Reversible Rhodopsin Dynamics in a Living Eye"

### I. Bleach Level Calculation

Early photointermediates, up to metarhodopsin I (Meta I), can absorb an additional photon and revert either back to rhodopsin or its isoform (9-cis-rhodopsin or isorhodopsin) in a process known as "photoreversal" (1, 2). Consequently, if a high-intensity flash is delivered before the majority of Meta I thermally converted to Metarhodopsin II (Meta II), a transient photo-equilibrium is established between rhodopsin and its early intermediates preceding Meta II (3).

Hence, considering the short-duration pulses used in this study, it is necessary to characterize how photoreversal affects the bleaching process. Following the flash bleaching model established by Gupta et.al. (4) and the isomerization diagram shown in Fig. 3A of the main text, the kinetic equations can be written as following:

$$R_0 \frac{dr(t)}{dt} = -\gamma_{RL} J_R(t) + \gamma_{LR} J_L(t) + \gamma_{M_1 R} J_{M_1}(t) \quad [1]$$

$$R_0 \frac{di(t)}{dt} = -\gamma_{IL} J_I(t) + \gamma_{LI} J_L(t) + \gamma_{M_1 I} J_{M_1}(t) \quad [2]$$

$$R_0 \frac{dl(t)}{dt} = -(\gamma_{LR} + \gamma_{LI}) J_L(t) - K_{LM_1} L(t) + \gamma_{RL} J_R(t) + \gamma_{IL} J_I(t) \quad [3]$$

$$R_0 \frac{dm_1(t)}{dt} = -(\gamma_{M_1 R} + \gamma_{M_1 I}) J_{M_1}(t) + K_{LM_1} L(t) - K_{M_1 M_2} M_1(t) \quad [4]$$

$$R_0 \frac{dm_2(t)}{dt} = K_{M_1 M_2} M_1(t) \quad [5]$$

where $R_0$ (in chromophore·µm$^{-3}$) denotes the concentration of rhodopsin in the dark state; $r(t) = R(t)/R_0$, $i(t) = I(t)/R_0$, $l(t) = L(t)/R_0$, $m_1(t) = M_1(t)/R_0$, $m_2(t) = M_2(t)/R_0$ are relative concentrations of rhodopsin, isorhodopsin, lumirhodopsin, Meta I and Meta II, respectively; $\gamma_{XY}$ denotes the quantum efficiency of the photically driven conversion from molecule $X$ to $Y$, $K_{XY}$ represents the thermal reaction rate constant (in s$^{-1}$) from $X$ to $Y$, and $J_X(t)$ represents the absorption rate (in photons·µm$^{-3}$·s$^{-1}$) of the species $X$ at time $t$ given by

$$J_X(t) = \int_0^{+\infty} J_X(\lambda, t)\, d\lambda \quad [6]$$

where $J_X(\lambda, t)$ is the spectral absorption rate of the species $X$ and is given by (4, 5):

$$J_X(\lambda, t) = \frac{\alpha_X(\lambda) X(t) I_{\mathrm{ROS}}(\lambda, t)}{H} \left(1 - e^{-H}\right) \quad [7]$$

where $\alpha_X(\lambda)$ (in µm$^2$·chromophore$^{-1}$) denotes the extinction coefficient of $X$ at wavelength $\lambda$, $I_{\mathrm{ROS}}$ (in photons·µm$^{-2}$·nm$^{-1}$·s$^{-1}$) represents the irradiance spectrum of the photon influx at the level of rod outer segments (ROS), and

$$H = \left[\alpha_R(\lambda) r(t) + \alpha_I(\lambda) i(t) + \alpha_L(\lambda) l(t) + \alpha_{M_1}(\lambda) m_1(t)\right] R_0 d \quad [8]$$

with $d$ being the length of ROS. $I_{\mathrm{ROS}}$ can be derived from experimentally measured $I_{in}$ at the cornea by (6)

$$I_{\mathrm{ROS}}(\lambda, t) = f_{wg} \tau I_{in}(\lambda, t) \quad [9]$$

where $f_{wg}$ represents the factor by which the rod waveguide condenses light into the ROS and $\tau$ is transmissivity of the ocular media. The extinction coefficients can be calculated as:

$$\alpha_X(\lambda) = \alpha_{X,\max} S_X(\lambda) \quad [10]$$

where $\alpha_{X,\max}$ is the peak extinction coefficient and $S_X$ is the normalized absorption spectrum, which can be calculated using the Lamb template function given as (7):

$$S_X(\hat{\lambda}) = \left[e^{a_1(b_1 - \hat{\lambda})} + e^{a_2(b_2 - \hat{\lambda})} + e^{a_3(b_3 - \hat{\lambda})} + c\right]^{-1} \quad [11]$$

where $\hat{\lambda} = \lambda_{X,\max}/\lambda$ with $\lambda_{X,\max}$ being the wavelength corresponding to the peak extinction coefficient. The template parameters are given in Table. S1. Substituting equations [9] and [10] into equations [7] and [8], and after some algebraic rearrangements, we obtain

$$\frac{J_X(\lambda, t)}{R_0} = (f_{wg} \tau \alpha_{R,\max} \gamma_{RL}) \cdot \frac{\alpha_{X,\max}}{\alpha_{R,\max}} \frac{S_X(\lambda) x(t) I_{in}(\lambda, t)}{\gamma_{RL} H} \left(1 - e^{-H}\right) \quad [12]$$

where $H$ becomes

$$H = \left[S_R(\lambda) r(t) + \frac{\alpha_{I,\max}}{\alpha_{R,\max}} S_I(\lambda) i(t) + \frac{\alpha_{L,\max}}{\alpha_{R,\max}} S_L(\lambda) l(t) + \frac{\alpha_{M1,\max}}{\alpha_{R,\max}} S_{M_1}(\lambda) m_1(t)\right] \alpha_{R,\max} R_0 d \quad [13]$$

with $1/(f_{wg}\tau\alpha_{R,\max}\gamma_{RL}) = 6.2 \times 10^7$ μm$^{-2}$ (8, 9), and $\alpha_{R,\max}R_0 d = 0.806$ (9).

The dynamics of relative concentrations for all photointermediates can be determined by solving the system of kinetic equations numerically (e.g., the Euler's method) based on equations [1] - [5] and [12] - [13], as well as the initial condition of $r(0) = 1$, $i(0) = l(0) = m_1(0) = m_2(0) = 0$. The values of parameters involved in the calculation are listed in Table. S1. The bleach level is then evaluated as the sum of the relative concentrations of lumirhodopsin, Meta I and Meta II at the end of the visual stimulus.

### II. Computational Model of ROS Contraction

We modeled the rapid contraction of ROS based on the voltage-dependent membrane tension model, which was previously developed to explain similar observations in the cone outer segment (COS) (10, 11). In brief, charge shift across the membrane during the R2 phase of the early receptor potential (ERP) leads to hyperpolarization of the disc membrane. Increased concentration of charges in the Debye layer increases their lateral repulsion on both sides of the membrane (12–14). Considering the volume conservation during the millisecond-scale dynamics, lateral stretching of the discs results in their axial contraction. The cumulative contraction of individual discs leads to the contraction of the photoreceptor OS by tens or even hundreds of nanometers.

**Quasistatic membrane model.** Earlier study of the rat rods has reported that a full bleach leads to a shift of $2 \times 10^5$ electronic charges during the R2 phase of the ERP (15). Considering that 2% of a total of $7 \times 10^7$ rhodopsins are embedded in the rod plasma membrane and nascent basal discs (16, 17), the charge shift per photoisomerization is estimated to be $q = 0.14$ electrons.

The transmembrane voltage change in the disc can be calculated by

$$\Delta V_m(t) = m_2(t) \cdot \frac{\sigma_{\text{rho}} \cdot q}{c_m} \quad [14]$$

where $m_2$ is the relative concentration of Meta II calculated in the previous section. $\sigma_{\text{rho}} = 23000$ molecules·μm$^{-2}$ is the area density of rhodopsin molecules in the disc membrane (17). $c_m = 1$ μF·cm$^{-2}$ is the specific membrane capacitance (15).

The charge displacement during ERP increases the repulsive forces of ions $\tau_e$, on the order of 0.1 $\text{mN} \cdot \text{m}^{-1} \cdot \text{V}^{-1}$ (10, 12). In the quasistatic model, increase in repulsive force is immediately balanced by the tension $\tilde{\tau}$ caused by the mechanical stretch: $\tilde{\tau}(t) = \tau_e(t)$. The associated normalized area expansion of a lipid membrane was derived as (10, 18)

$$\frac{\Delta A(t)}{A_0} = \frac{1}{\alpha} \ln\left[1 + \frac{\tilde{\tau}(t)}{\beta}\right] \quad [15]$$

where $A_0$ denotes the resting area of a disc membrane patch (2.1 μm$^2$ in rat rods (19)), $\Delta A$ correspond to the tension-dependent area expansion of the disc membrane. $\alpha = 8\pi\kappa_c/k_B T$ and $\beta = \pi^2\kappa_c/A_0$, where $\kappa_c$ is the bending modulus, $k_B$ is the Boltzmann's constant, $T = 310$ $K$ is the absolute temperature.

Considering the volume conservation within each disc during the millisecond-scale dynamics, the associated axial contraction of each disc can be calculated by

$$\Delta h(t) = -\frac{\Delta A(t)}{A_0} \cdot h_0 \quad [16]$$

where $h_0 = 18$ nm is the height (thickness) of individual discs (20).

The observed contraction in ROS is the cumulative contribution of individual discs,

$$\Delta\text{OPL}(t) = \Delta h(t) \cdot N_{\text{disc}} \cdot n_{\text{index}} \quad [17]$$

where $N_{\text{disc}} = 750$ is the number of discs per ROS (19, 20). $n_{\text{index}} = 1.41$ is the average refractive index of the ROS (21).

When neglecting passive discharge of the hyperpolarized membrane, the maximum change in the transmembrane voltage can be calculated by replacing $m_2(t)$ in equation [14] by the bleach level, denoted as BL:

$$\Delta V_{m,\ \max} = \text{BL} \cdot \frac{\sigma_{\text{rho}} \cdot q}{c_m} \quad [18]$$

The area expansion of individual discs and the amplitude of ROS contraction can be calculated based on equations [15]-[17]. Fig. S1 illustrates the overall good fit between the measured rapid contraction amplitude (colored dots) and the contraction amplitude calculated based on the voltage-dependent membrane tension model (gray curve). At the best fit, we obtained an initial tension $\tau_0$ of 0.59 μN·m$^{-1}$ and a bending modulus $\kappa_c = 0.23 \times 10^{-19}$ N · m. The typical ranges of the membrane surface tension and bending modulus in the literature are $0.1 - 1$ μN·m$^{-1}$ and $0.5 - 2 \times 10^{-19}$ N · m (10), respectively, pretty close to our fitted values.

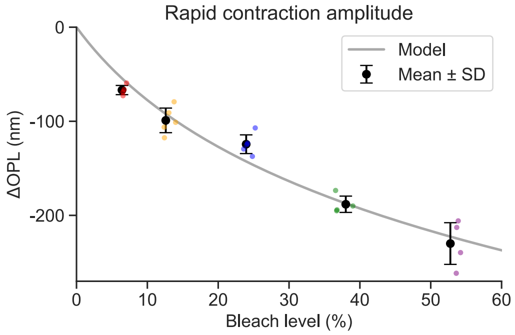


**Fig. S1.** The measured contraction amplitude (colored dots) fits the prediction from the voltage-dependent membrane tension model (gray curve).

**The planar friction model (PF model).** Although contraction amplitude calculated based on the voltage-dependent membrane tension model aligned well with experimental measurements, the quasistatic model failed to capture the dynamics of the ROS contraction (see blue curves in Figs. 4B-C). Much slower contraction process compared to the driving force $\tau_e(t)$ indicates the necessity of incorporating viscoelastic effects.

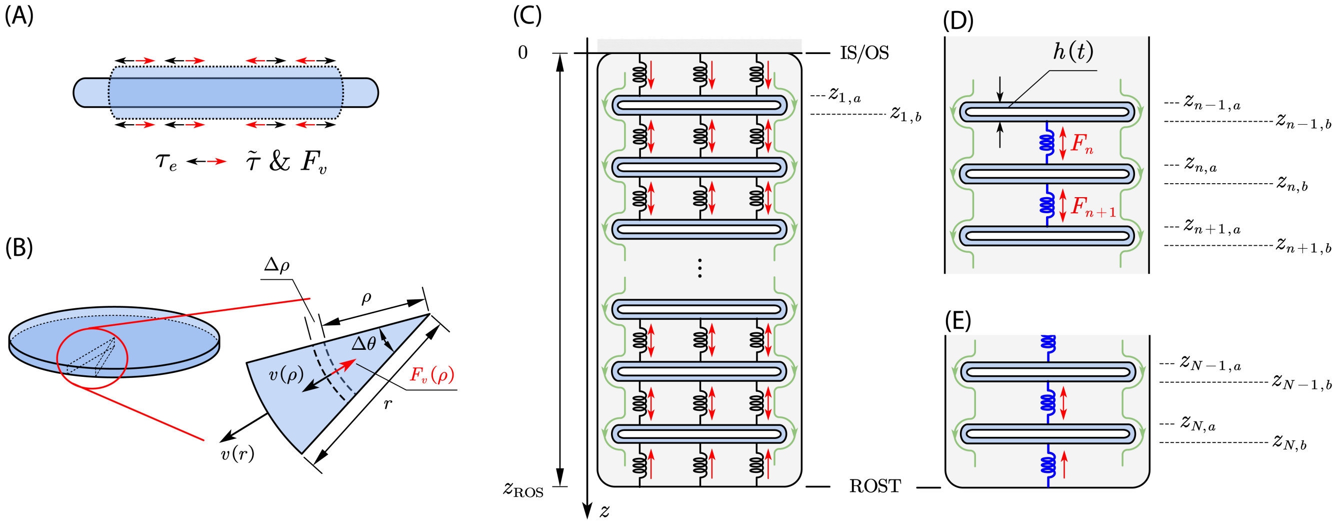


**Fig. S2.** Viscoelastic modeling of the ROS contraction. (A) Axial contraction of a single disc due to increased lateral repulsive force and conservation of the disc volume. $\tau_e$: the tension caused by repulsive forces of ions, which is related to the transmembrane voltage. $\tilde{\tau}$: the lipid membrane tension. $F_v$: viscous force due to the medium. (B) Geometry of the radially expanding disc. The elemental area of interest is the differential patch on the arc sector from a disc between the two dashed lines, $[\rho, \rho + \Delta\rho] \times [0, \Delta\theta)$. (C) A schematic of ROS with a distributed spacer network. IS/OS junction is modeled as a fixed boundary and ROST as a free end. Both boundaries of each disc are indexed. Red arrows showing the direction of axial stretching of spring-like spacers due to axial contraction of discs. (D) A free-body diagram for an example disc in the middle of ROS. (E) A free-body diagram illustrating the ROST boundary.

As illustrated in Fig. S2A-B, we modeled the contraction of individual disc membranes by incorporating a linear viscous force $F_v$. The friction force on a differential patch element, induced by surrounding fluid can be calculated as (Fig. S2B),

$$F_v(\rho) = -\eta_m \cdot v(\rho) \cdot \rho\Delta\theta\Delta\rho \qquad [19]$$

where $\eta_m$ (in N·s·m$^{-3}$) is the damping coefficient, and $\rho\Delta\theta\Delta\rho$ is the area of the differential patch (Fig. S2B).

As derived in our previous publication, the power dissipated through the linear viscous force is (10),

$$P_v = \frac{\pi\eta_m r^2 \dot{r}^2}{2} \qquad [20]$$

The power done by the membrane during its expansion or contraction is (10),

$$P_m = -2\pi\tau_{\mathrm{total}} r\dot{r} \qquad [21]$$

where $\tau_{\text{total}} = \tilde{\tau} - \tau_e = \beta(e^{2\alpha x} - 1) - \tau_e$ is the total membrane tension considering both the exponential restoring force (see equation [15]) and the repulsive forces of ions. $x(t) = (r(t) - r_0)/r_0$, and $r_0$ is the initial radius of the disc patch.

Given that the work done by the membrane converts to the heat dissipation through viscosity, we have $P_v = P_m$, which can be simplified as,

$$\frac{\eta_m r_0^2}{4}\dot{x} + \beta\left(e^{2\alpha x} - 1\right) = \tau_e(t) \qquad [22]$$

The value of $x$ at rest status, denoted by $x_0$, can be calculated based on $\beta(e^{2\alpha x_0} - 1) = \tau_0$, where $\tau_0$ is the initial tension. Solving for $x(t)$ numerically allows the axial contraction for a single disc to be determined as

$$\Delta h(t) = -2\left[x(t) - x_0\right] \cdot h_0 \qquad [23]$$

By fitting the viscoelastic membrane model to the experimentally measured $\Delta$OPL (see black and orange traces in Fig. S3A), we obtained a damping coefficient of $1.22 \times 10^6$ N·s·m$^{-3}$, which corresponds to a linear damping behavior induced by a cytoplasm sheet with a thickness of $\sim$3 nm(22, 23).

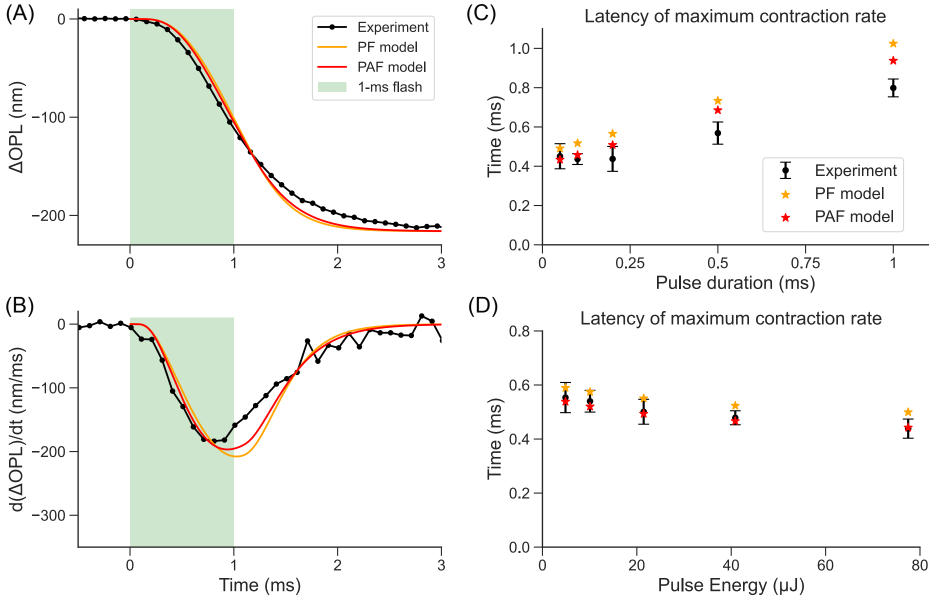


**Fig. S3.** ROS contraction measured and calculated based on viscoelastic models. (A) Experimentally measured $\Delta$OPL response to a 1-ms flash and predictions of the planar friction (PF) model and the planar & axial friction (PAF) model. (B) Contraction rate of the traces in (A). (C)-(D) Comparison of the latency predicted by viscoelastic models (orange and red stars) and measured in experiments (black error bars) with varying pulse durations and varying pulse energies, respectively.

Although the viscoelastic membrane model matches the experimental traces much better than a quasistatic model, it cannot precisely account for the latency of the maximum contraction rate across different flash configurations (see orange traces and stars in Figs. S3B-D).

**The planar & axial friction model (PAF model).** Another source of damping could arise from the friction when discs move axially within the viscous cytoplasm. Cryo-electron tomography has revealed protein spacers that act as springs and connect the adjacent discs (24). The shrinkage of each disc results in the stretching of the spacers (Fig. S2C, red arrows) which drive axially the discs and the ROS tip (ROST). As individual discs move along the axial direction, the viscous force induced by surrounding cytoplasm dampens the contraction process. We assume that

1. All spacers between two adjacent discs are represented as a single lumped spring with a spring constant $k_s$ (Fig. S2C-E), and this value is estimated to be 15.5 N·m$^{-1}$ (21).

2. The inner segment/outer segment junction (IS/OS) is fixed during this process, i.e., $z_{0,b} = 0$.

3. ROST moves together with the bottom surface of the $N^{\text{th}}$ disc, i.e., $z_{N+1,a} = z_{N,b}$.

4. The mass of individual discs is considered negligible in the model, meaning that the system is overdamped. This is supported by the fact that the maximum contraction rate observed in our measurements was approximately 200 nm/ms.

The corresponding kinetic energy of a disc moving at this speed is equivalent to the elastic energy of a spacer expanding by $3 \times 10^{-4}$ nm. In comparison, our modeled results show that the expansion of spacers reaches up to several 0.01 nm. Since the elastic energy of individual spacers is three orders of magnitude larger than the maximum kinetic energy of individual discs, we neglected the effect of mass in the PAF model.

Under these assumptions, for each spacer, we have

$$F_n = k_s(\Delta z_{n,a} - \Delta z_{n-1,b}), \quad \forall n \in [1, N+1] \tag{24}$$

**The volume conservation constraint**. The volume of the $n^{\text{th}}$ disc can be found as

$$(z_{n,b} - z_{n,a})\pi r_n^2 = V_0 \tag{25}$$

Taking the time derivative on both sides, we have

$$\frac{\dot{z}_{n,b} - \dot{z}_{n,a}}{z_{n,b} - z_{n,a}} = -\frac{2\dot{r}_n}{r_n} \tag{26}$$

Assume small deformations in discs and denote $\dot{h}_n = \dot{z}_{n,b} - \dot{z}_{n,a}$, then

$$\frac{\dot{h}_n}{h_0} = -\frac{2\dot{r}_n}{r_0} \tag{27}$$

where $h_0$ and $r_0$ denote the initial thickness and radius of all discs, respectively. Similarly, taking the total differential, we have

$$\frac{\Delta z_{n,b} - \Delta z_{n,a}}{h_0} = -\frac{2\Delta r_n}{r_0} \tag{28}$$

**The energy conservation constraint**. In the viscoelastic membrane model, we assumed that the power generated by the membrane tension is dissipated through viscosity when each disc membrane expands laterally. Here, we need to further consider the power exerted by the axial forces on discs, which can be written as

$$P_{\text{ax}} = F_{n+1}\dot{z}_{n,b} - F_n\dot{z}_{n,a} + f_n\frac{\dot{z}_{n,a} + \dot{z}_{n,b}}{2} \tag{29}$$

where $f_n$ denotes the axial friction force exerted on the side walls of the $n^{\text{th}}$ disc given as

$$\begin{aligned} f_n &\approx -\eta_v A_s \overline{\dot{z}_n} && [30] \\ &= -\eta_v(2\pi r_0 h_0)\frac{\dot{z}_{n,a} + \dot{z}_{n,b}}{2} \\ &= -\pi\eta_v r_0 h_0(\dot{z}_{n,a} + \dot{z}_{n,b}) && [31] \end{aligned}$$

with $\eta_v$ denoting the damping coefficient.

Assume the mass of discs is negligible, then $F_n = f_n + F_{n+1}$, hence

$$\begin{aligned} P_{\text{ax}} &= F_{n+1}\dot{z}_{n,b} - F_n\dot{z}_{n,a} + (F_n - F_{n+1})\frac{\dot{z}_{n,a} + \dot{z}_{n,b}}{2} && [32] \\ &= (F_n + F_{n+1})\frac{\dot{z}_{n,b} - \dot{z}_{n,a}}{2} && [33] \\ &= -(F_n + F_{n+1})\frac{h_0}{r_0}\dot{r}_n && [34] \end{aligned}$$

Combining the lateral power derived from the previous viscoelastic membrane model (see Eqs. [20] and [21]) and considering two membrane surfaces (top and bottom), we obtain

$$-(F_n + F_{n+1})\frac{h_0}{r_0}\dot{r}_n - 4\pi\tau_{\text{total}} r_0\dot{r}_n = \pi\eta_m r_0^2\dot{r}_n^2 \tag{35}$$

Dividing both sides by $\dot{r}_n$ and substituting the equation [27]:

$$(F_n + F_{n+1})\frac{h_0}{r_0} + 4\pi\tau_{\text{total}} r_0 = \frac{\pi\eta_m r_0^3\dot{h}_n}{2h_0} \tag{36}$$

After some algebraic rearrangements, we obtain

$$\dot{h}_n = \dot{z}_{n,b} - \dot{z}_{n,a} = \frac{2h_0^2}{\pi\eta_m r_0^4}(F_n + F_{n+1}) + \frac{8h_0}{\eta_m r_0^2}\left[\beta\left(e^{-\alpha(\Delta h_0 + \Delta z_{n,b} - \Delta z_{n,a})/h_0} - 1\right) - \tau_e\right] \tag{37}$$

where $\Delta h_0$ is the axial expansion of each disc in the resting status, which can be calculated based on $\beta(e^{-\alpha\Delta h_0/h_0} - 1) = \tau_0$, where $\tau_0$ is the initial tension.

In summary, the complete dynamic system combines equations [24], [30] and [37]:

$$\begin{cases} F_n = k_s(\Delta z_{n,a} - \Delta z_{n-1,b}), \quad \forall n \in [1, N+1] \\ \dot{z}_{n,b} + \dot{z}_{n,a} = \dfrac{1}{\pi\eta_v r_0 h_0}(F_{n+1} - F_n), \quad \forall n \in [1, N] \\ \dot{z}_{n,b} - \dot{z}_{n,a} = \dfrac{2h_0^2}{\pi\eta_m r_0^4}(F_n + F_{n+1}) + \dfrac{8h_0}{\eta_m r_0^2}\left[\beta\left(e^{-\alpha(\Delta h_0 + \Delta z_{n,b} - \Delta z_{n,a})/h_0} - 1\right) - \tau_e\right], \quad \forall n \in [1, N] \\ z_{0,b} = 0 \\ z_{N+1,a} = z_{N,b} \end{cases} \tag{38}$$

By fitting the PAF model to the experimentally measured ΔOPL (see black and red traces in Figs. S3A)., we obtained damping coefficients of $4.0 \times 10^5$ N·s·m$^{-3}$ for the lateral expansion of disc membranes and $4.4 \times 10^4$ N·s·m$^{-3}$ for the axial movement of each disc. These damping coefficients correspond to the linear damping behavior caused by cytoplasmic sheets with thicknesses of approximately 10 nm and 100 nm, respectively (22).

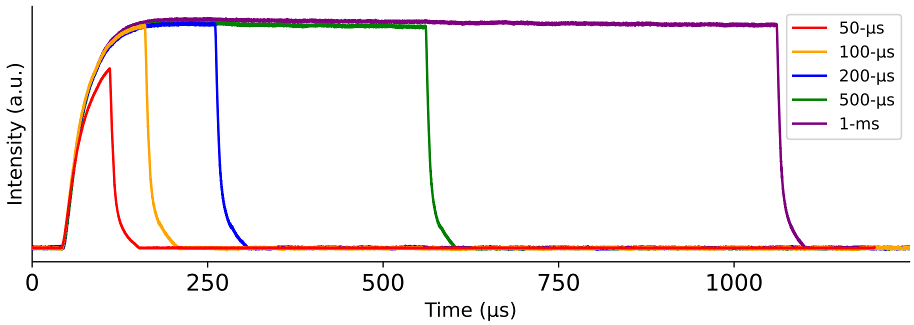


**Fig. S4.** Temporal profiles of green flashes (520 nm) with varying pulse durations.

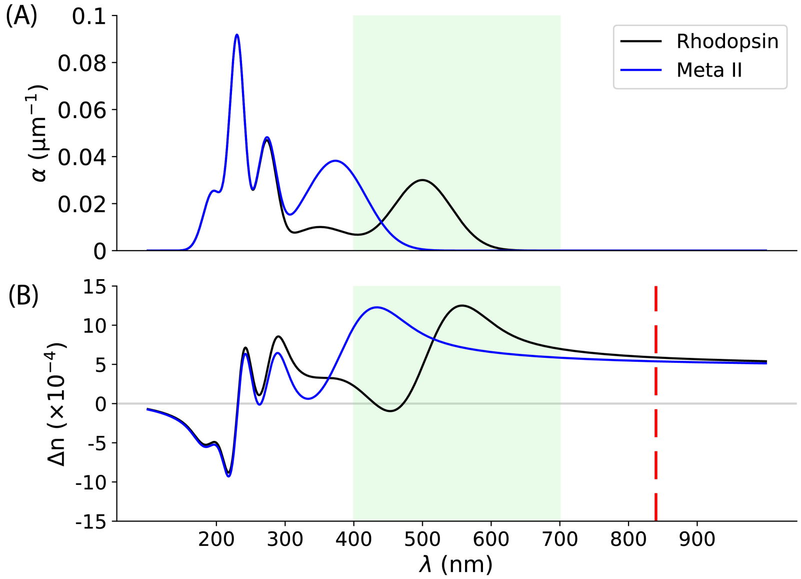


**Fig. S5.** Refractive index variation calculated based on the Kramers-Kronig transform. (A) Absorption spectrum (black) of an idealized pigment mimicking rhodopsin (25). The Meta II absorption spectrum (blue) constructed by shifting the absoption band centered at 500 nm to 380 nm. (B) Refractive index changes derived by the Kramers-Kronigs relationship using absorption spectra in the top panel. Vertical line at OCT center wavelength (840 nm) intercepts the $\Delta$n curves at $5.88 \times 10^{-4}$ and $5.39 \times 10^{-4}$ for rhodopsin and Meta II, respectively.

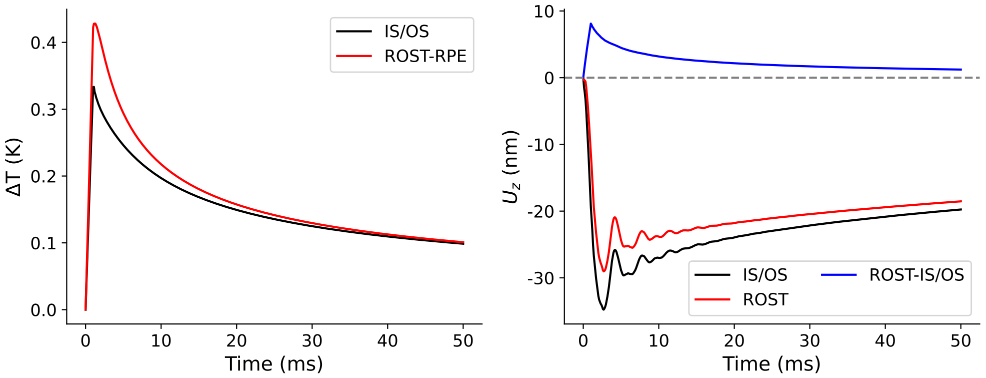


**Fig. S6.** Thermo-mechanical simulation of the retinal deformation. Simulation parameters: ocular transmittance = 60%, pulse energy (measured in front of the cornea) = 450 µJ, pulse duration = 1 ms, illumination area = 1.88 $mm^2$. Left: simulated temperature change of various planes at the beam center. Right: simulated axial displacements of various planes at the beam center. Blue trace represents the change in the physical length of the rod outer segment. The ripples are due to mechanical waves reflected off the boundary.

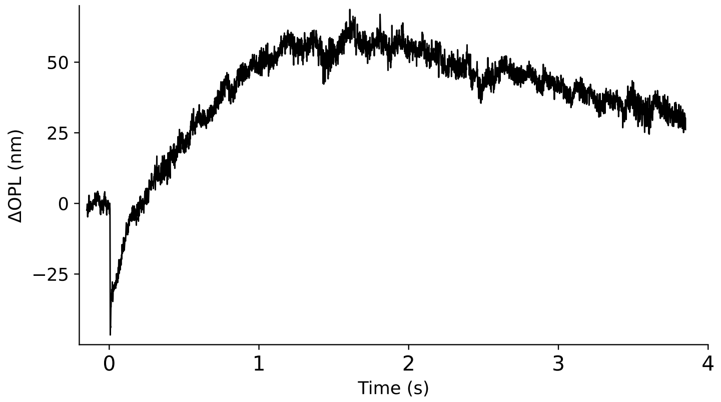


**Fig. S7.** An example ORG response evoked by a single green flash (2.4 µJ, 100 µs) recorded at 1 kHz for 4 seconds, showing the recovery from the rapid contraction.

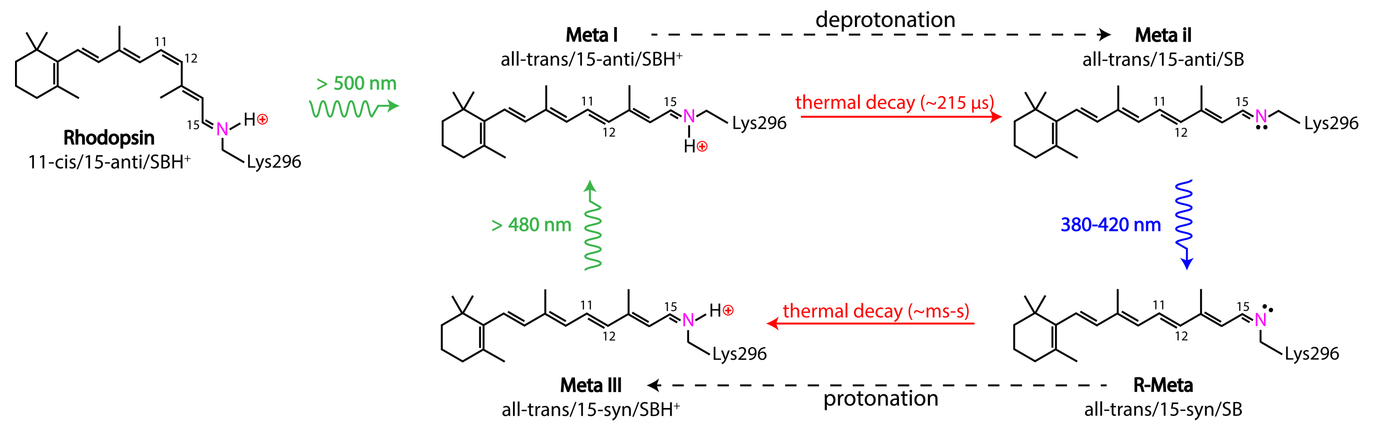


**Fig. S8.** Photointermediate transition pathways under alternating green and UV flashes.

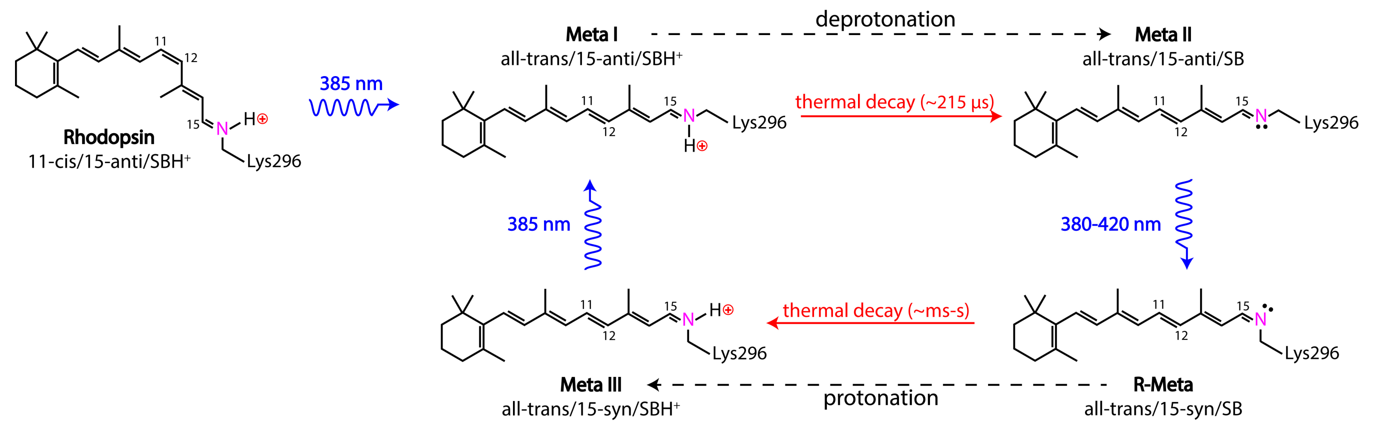


**Fig. S9.** Photointermediate transition pathways under UV flashes.

**Table S1.** Parameters used for modeling flash photolysis in rats

| Quantum efficiency | Value |
|---|---|
| $\gamma_{RL}$ (26) | 0.67 |
| $\gamma_{IL}$ (26) | 0.23 |
| $\gamma_{LR}$ (27) | 0.42 |
| $\gamma_{LI}$ (27) | 0.07 |
| $\gamma_{M_1R}$ (28) | 0.33 |
| $\gamma_{M_1I}$ (28) | 0.06 |
| **Thermal rate constant ($s^{-1}$) (16)** | **Value** |
| $K_{LM_1}$ | $2.17\times10^4$ |
| $K_{M_1M_2}$ | $4.65\times10^3$ |
| **Relative peak extinction coeff. (4)** | **Value** |
| $\alpha_{I,\text{max}}$ / $\alpha_{R,\text{max}}$ | 1.13 |
| $\alpha_{L,\text{max}}$ / $\alpha_{R,\text{max}}$ | 1.18 |
| $\alpha_{M_1,\text{max}}$ / $\alpha_{R,\text{max}}$ | 1.06 |
| **Template params. for $S_X$ (7)** | **Value** |
| $(a_1, b_1)$ | (70, 0.880) |
| $(a_2, b_2)$ | (28.5, 0.924) |
| $(a_3, b_3)$ | (-14.1, 1.104) |
| $c$ | 0.655 |
| **Wavelength at peak absorption (nm) (29)** | **Value** |
| $\lambda_R$ | 498 |
| $\lambda_I$ | 486 |
| $\lambda_L$ | 492 |
| $\lambda_{M_1}$ | 478 |
